# Supernumerary proteins of the human mitochondrial ribosomal small subunit are integral for assembly and translation

**DOI:** 10.1101/2022.06.15.495910

**Authors:** Taru Hilander, Geoffray Monteuuis, Ryan Awadhpersad, Krystyna L. Broda, Max Pohjanpelto, Elizabeth Pyman, Sachin Kumar Singh, Tuula A. Nyman, Isabelle Crevel, Robert W. Taylor, Ann Saada, Diego Balboa, Brendan J. Battersby, Christopher B. Jackson, Christopher J. Carroll

## Abstract

Mitochondrial ribosomes (mitoribosomes) have undergone substantial structural remodelling throughout evolution. Compared to their prokaryotic counterparts, mitoribosomes show a substantial loss of ribosomal RNA, whilst acquiring unique protein subunits located on the periphery of the ribosomal subunit structures. We set out to investigate the functional properties of all 14 unique (mitochondrial-specific or supernumerary) human mitoribosomal proteins in the small subunit. Using genome editing with CRISPR-Cas9, we made knockouts for each subunit in HEK293 cells to study the effect on mitoribosome assembly and function in protein synthesis. Unexpectedly, we show that each supernumerary knockout leads to a unique mitoribosome assembly defect with variable impact on mitochondrial protein synthesis. Our data demonstrates that all supernumerary subunits are essential structural components except mS37. Surprisingly, we found the stability of mS37 was reduced in all our supernumerary knockouts of the small and large ribosomal subunits as well as patient-derived lines with mitoribosome assembly defects. We identified that a redox regulated CX_9_C motif in mS37 was essential for protein stability, suggesting a potential mechanism to regulate mitochondrial protein synthesis. Together, our findings support a modular assembly of the human mitochondrial small ribosomal subunit mediated by essential supernumerary subunits and identify a redox regulatory role involving mS37 in mitochondrial protein synthesis in health and disease.

## Introduction

Mitochondria are cellular organelles of prokaryotic origin where ATP is synthesized by oxidative phosphorylation (OXPHOS). Whilst most mitochondrial genes have been transferred to the nucleus during evolution, human mitochondria continue to harbour their own compacted circular genome. Human mitochondrial DNA (mtDNA) encodes 13 highly hydrophobic structural subunits of the OXPHOS enzyme complexes, while the remaining genes encode 22 tRNAs and 2 rRNAs required for mitochondrial protein synthesis (Amunts et al., 2015; Desai et al., 2017, 2020; Greber et al., 2015; Itoh et al., 2021; Koripella et al., 2020; Kummer and Ban, 2021; Silva et al., 2015).

The mature human mitoribosome (55S) is composed of a small (28S, mtSSU) and a large (39S, mtLSU) subunit. The human mtSSU is composed of 30 proteins and a 12S rRNA, while the mtLSU consists of 52 proteins, a 16S rRNA and a copy of mitochondrial valine transfer RNA (Brown et al., 2014). Across eukaryotic taxa, mitoribosomes differ in size and the protein to RNA ratio, including mammals (55S), budding yeast (74S) and plants (78S) (Figure S1) (Amunts et al., 2015; Desai et al., 2017; Greber et al., 2015; Saurer et al., 2019; Waltz et al., 2020). The increase in mitoribosomal protein abundance derives from increased extensions in shared homologous proteins and the addition of mitochondrial-specific proteins, also known as supernumerary mitochondrial ribosomal proteins (snMRPs) (Rackham and Filipovska, 2014). The recent cryo-EM structures of mitoribosomes from human, porcine and yeast reveal that the supernumerary proteins are extensions occupying completely new positions, rather than a replacement of lost rRNA segments (Amunts et al., 2015; Greber et al., 2015; Kummer and Ban, 2021). It is currently unclear whether these snMRPs primarily compensate for the reduced rRNA abundance or participate in additional functions important for mitochondrial protein synthesis. With the growing number of reports of human inherited diseases with mitochondrial protein synthesis defects and tissue-specific symptoms, including several mitoribosomal subunit proteins (Table S1), it is important to understand the role of these snMRPs.

The aim of our study was to determine the role of snMRPs for assembly of the human mtSSU and function in mitochondrial protein synthesis. We used genome editing to generate knockout cell lines for the snMRPs of the human mtSSU. Collectively, we find distinct phenotypes for each of the snMRPs on mitoribosome assembly and protein synthesis, and identify a novel role for mS37 in regulation of mitochondrial protein synthesis.

## Results

### CRISPR-mediated knockout of snMRPs of the mtSSU

We used CRISPR-Cas9 to generate knockout cell lines for each of the human snMRPs in HEK293. The nomenclature for mitoribosomal subunit proteins that has been adopted in several recent studies was applied here (Figure 1A) (Ban et al., 2014; Greber et al., 2015; Kummer and Ban, 2021). Since snMRPs are distributed on the periphery of the mtSSU and mtLSU, positions not previously occupied by the structural rRNA present in the bacterial ribosome, these proteins could have acquired additional mitochondrial-specific interactions for protein synthesis (Figure S2). We designed two guide RNAs (gRNAs) to all supernumerary proteins in the mtSSU: mS22, mS23, mS25, mS26, mS27, mS29, mS31, mS33, mS34, mS35, mS37, mS38, mS39 and mS40. The strategy was to excise a region encompassing the start codon, whilst ensuring no alternative initiation codons were present (Table S2). We successfully obtained clonal knockout cell lines for all 14 snMRPs of the mtSSU (Figure S3). All snMRP^KO^ cells were supplemented with uridine and pyruvate to bypass any potential growth defect caused by respiratory chain deficiency (Hart et al., 2015). Immunoblotting confirmed complete knockout in all snMRP^KO^ cells and revealed that each knockout had variable effects on the abundance of other mtSSU subunits (Figure 1B). Unfortunately, commercial antibodies were unavailable for mS31, mS33 and mS38 but the knockout allele was confirmed by PCR (Figure S3). In contrast to other mtSSU proteins, mS37 was uniformly decreased in all snMRP^KO^ cell lines, suggesting that amongst mtSSU proteins, mS37 protein is uniquely regulated.

**Figure 1.**
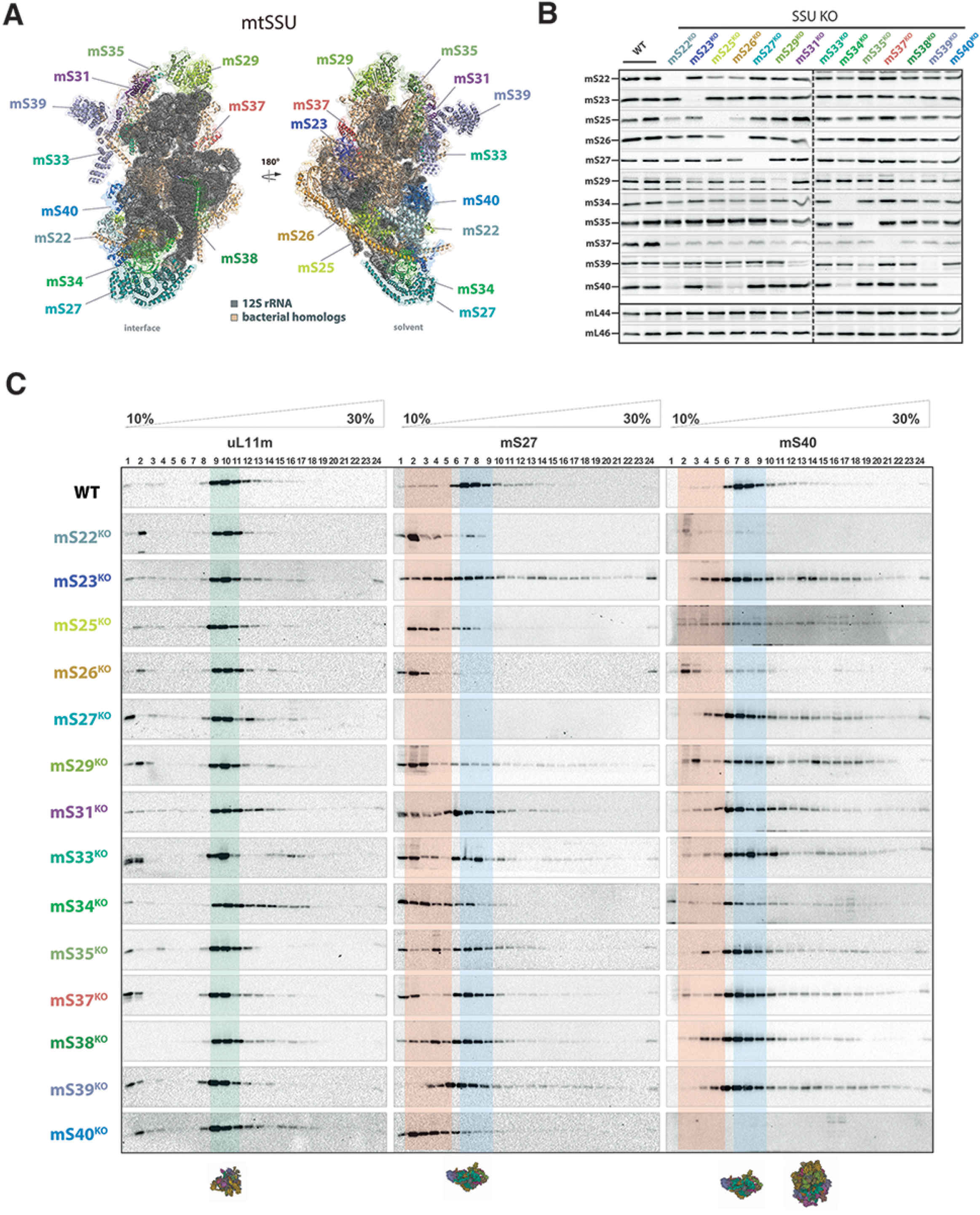
CRISPR-Cas9 mediated knockout of supernumerary proteins (snMRPs) of the human small mitoribosomal subunit (mtSSU) and effect on mitoribosome assembly. **(A)** snMRPs of the mtSSU (28S). Denomination of cryo-EM resolved mammalian mtSSU structure (PDB: 5AJ3) highlighting snMRPs of the mtSSU. Bacterial homologs in orange, 12S RNA in dark grey. **(B)** Immunoblot confirmation of knockout in HEK293 cells. The dotted lines indicate the border of separate immunoblots. **(C)** Isokinetic sucrose gradients of the 14 snMRP knockouts reveals mtSSU assembly defects. Fully assembled mtLSU fractions (green); fully assembled mtSSU fractions 6-8 (blue), intermediates and loss of assembly (orange).

Next, we assessed the status of assembled mitoribosomal subunits in snMRP^KO^ cell lines. Isokinetic sucrose gradients revealed a unique pattern of mtSSU assembly defects for each of the snMRP^KO^ cell lines (Figure 1C), using antibodies against early (mS27) and intermediate (mS40) assembling MRPs (Figure S4). The stability of the small mitoribosomal proteins mS27 and mS40 have been identified as markers for the step-wise assembly from the early to intermediate stage, respectively (Bogenhagen et al., 2018). In our snMRP^KO^ cell lines, we observed a shift towards an intermediate stage was most prominent in mS22^KO^, mS26^KO^, mS34^KO^ and mS40^KO^ cell lines. Assembly of the mtLSU was unaffected in our snMRP knockout cell lines (Figure 1C). Whether assembly intermediates lacking individual subunit proteins would be functional in mitochondrial protein synthesis is not known.

### Quantitative label-free proteomics reveals the impact on abundance of mtSSU proteins in snMRP knockouts

We performed quantitative label-free LC-MS/MS proteomic analysis for our snMRP^KO^ cell lines to assess quantitatively the abundance of mtSSU proteins. We visualised the fold-changes for all 14 snMRP^KO^ cell lines by mapping of the steady-state level with a 1.5-fold-expression change cut-off to the structure of the assembled 55S mitoribosome (Figure 2). Each snMRP^KO^ had a unique impact on the stability of other mtSSU proteins, and largely reflected the outcome for the assembled mtSSU observed in isokinetic sucrose gradients (Figure 1C). However, the mS37^KO^ cell line did not reveal any significant change in abundance of mtSSU proteins, corroborating the observation from the sucrose gradients. This suggests that the abundance of mS37 does not affect mtSSU assembly. Importantly, the timing of protein incorporation during mtSSU assembly (Bogenhagen et al., 2018) does not appear to be a reliable predictor for the steady-state abundance of the mtSSU. For example, mS25 and mS26 are known to be incorporated in the late stages of mtSSU assembly (Figure S4) and yet, our knockout cell lines for these subunits had the most deleterious impact on the mtSSU abundance. Interestingly, even proteins that do not bind the 12S rRNA, such as mS22, can generate a profound assembly defect for the mtSSU. Together, this data shows that knockout of snMRPs cause mtSSU assembly defects which are unique in each individual snMRP^KO^ cell line.

**Figure 2.**
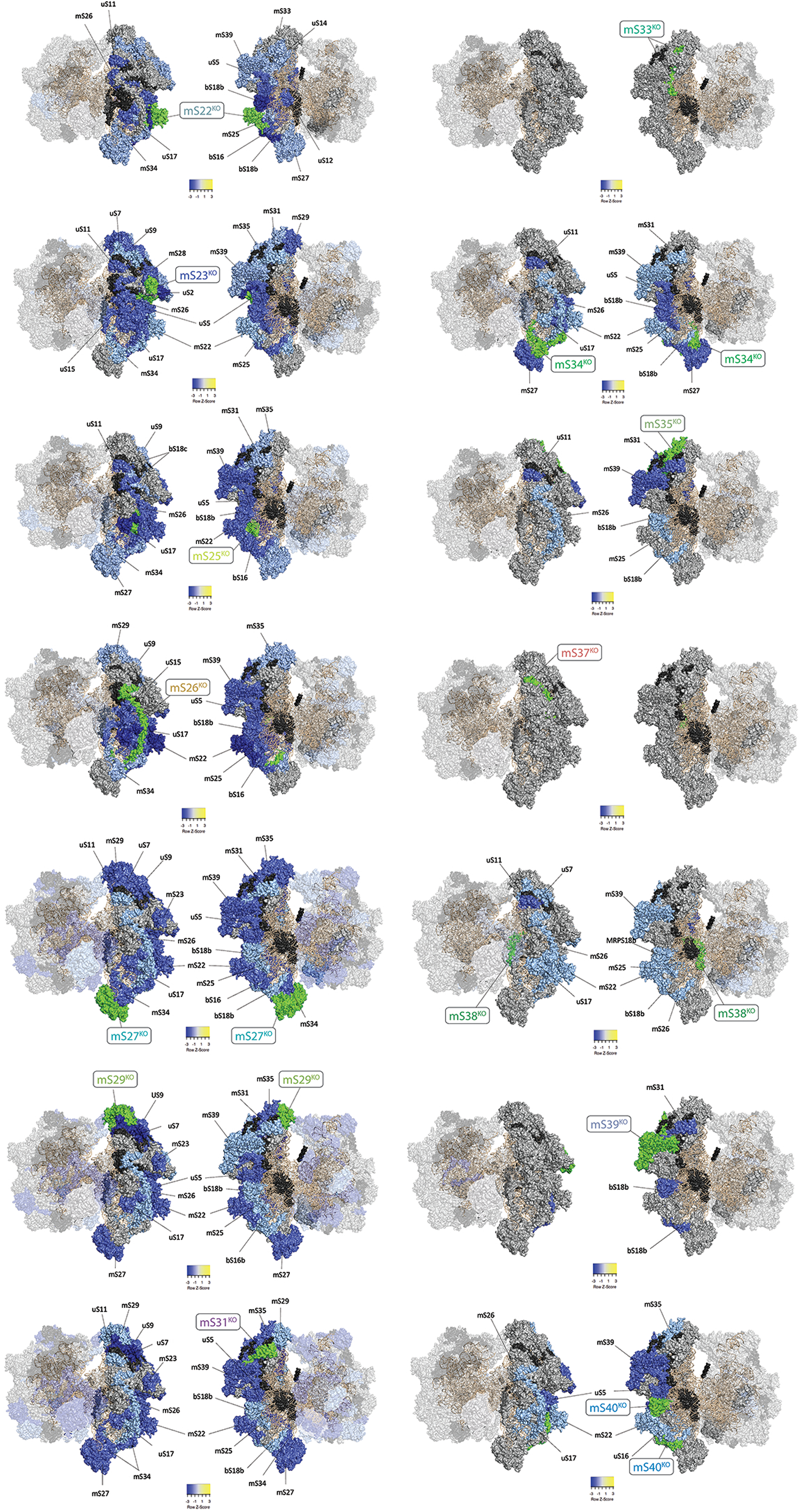
Proteomic expression mapping of the snMRP^KOs^ onto 55S mitoribosome structure. Ribosome structures based on (PDB entry: 5AJ4) and visualised using Pymol 2.7.1. Proteomic cut-off threshold for visualisation was 1.5-fold change from wild-type. For each snMRP^KO^ two images shown rotated 180°including both mtSSU and mtLSU. The snMRP^KO^ cell line is indicated in green and labelled in rectangular box. Proteins with differential expression are labelled. A gradient of blue/yellow colours indicates level of decrease/increase in expression (scale shown). Ribosomal RNA is orange. Proteins not changed in expression in grey. Proteins not reliably measured in black.

### Hierarchical clustering of mtSSU protein expression reveals assembly modules

While the structure of the human mitochondrial ribosome has been elucidated at high-resolution, far less is known about its assembly pathway. A recent study determined the kinetics by which ribosomal proteins are incorporated into the mtSSU (Figure S4), predicting that clusters of neighbouring proteins were first assembled into modules prior to incorporation into the growing complex (Bogenhagen et al., 2018). The specific proteins with reduced expression observed in each of our snMRP^KO^ cell lines are neighbouring proteins with close interactions in the mtSSU (Figure 2). Therefore, our data could enable predictions on the composition of the modular units that are assembled into the mtSSU.

We used the fold-expression changes of 23 out of the 30 mtSSU proteins that were reproducibly quantified in at least 10 of the cell lines for hierarchical clustering analysis (Supplementary Data File 1) (Figure 3A). We assessed whether the assigned clusters could be inferred as modules by comparing to the positions of ribosomal proteins in the mtSSU structure (Figure 3B). With remarkably few exceptions, hierarchical clustering accurately assigned mtSSU proteins to two broad groups, (1) head and platform and (2) body and foot. Furthermore, the predicted modules consisted of neighbouring proteins with surface interactions (Amunts et al., 2015), demonstrating the predictive power of the hierarchical clustering data for identifying interacting proteins, and indicating that for each module the constituent proteins were highly reliant on each other for their stability.

**Figure 3.**
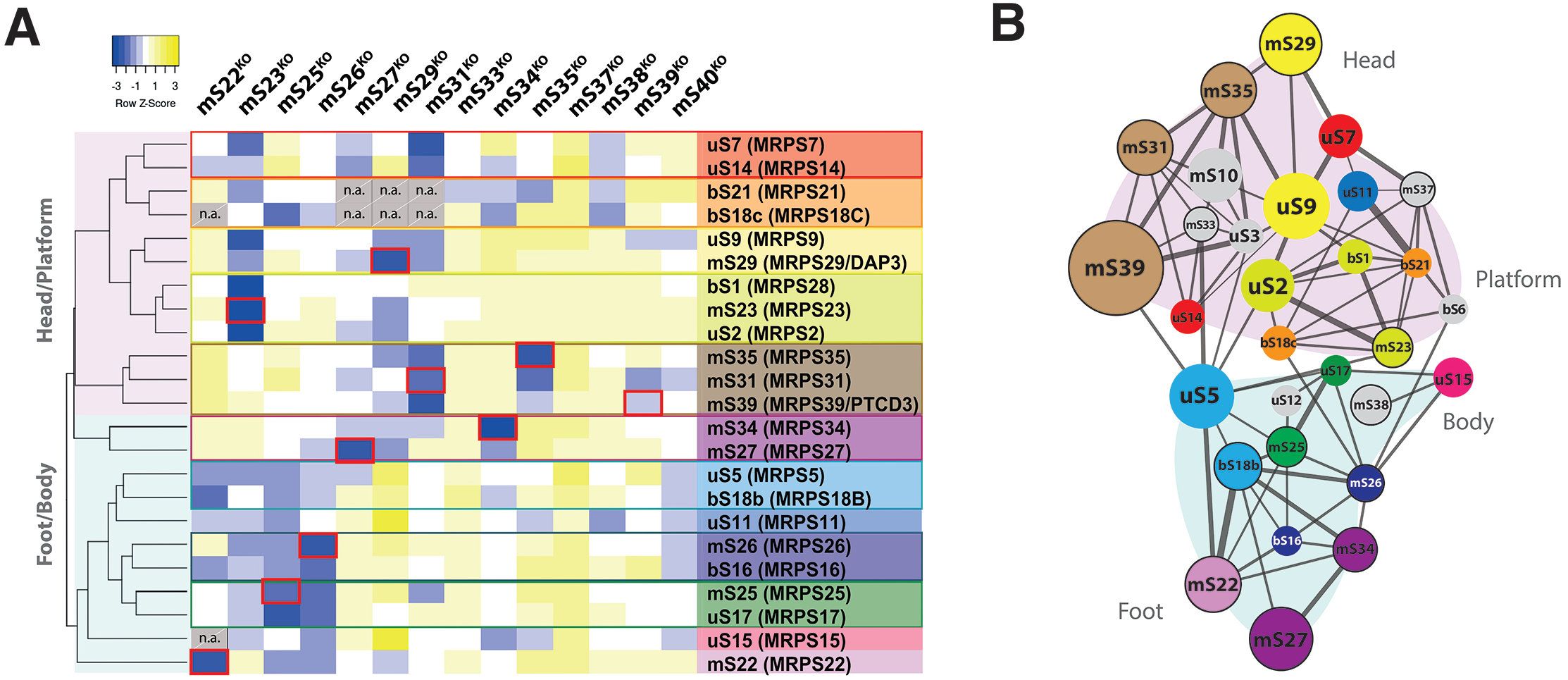
Hierarchical clustering of mtSSU protein expression reveals the composition of assembly modules of the mtSSU. **(A)** Hierarchical clustering of quantitative label-free proteomic data of mtSSU protein expression in snMRP knockouts compared to wild-type. A gradient of blue/yellow colours indicates level of decrease/increase in expression (scale shown). NA in grey boxes indicates data not available. Proteins with highest similarity in expression according to their clustering in the dendrogram are depicted with the same colour. Red outline in heatmap indicates respective knockout. (B) Surface interaction map of mtSSU proteins (adapted from (Amunts et al., 2015)) with superimposed colour code from protein expression from left panel. Line thickness indicates shared surface. Size of spot indicates protein size. Black circle indicates snMRPs.

We next assessed whether the expression levels of known mtSSU assembly factors correlates with assembly status of the mtSSU and therefore could allow for potential discovery of novel mtSSU assembly factors. The detected assembly factors were indeed altered in response to specific snMRP knockout (Figure S5).

### Knockout of mtSSU snMRPs causes distinct translation defects and OXPHOS dysfunction

To address the functional consequences of snMRP deficiency, we used metabolic labelling of ^35^S-methionine/cysteine to analyse mitochondrial protein synthesis. After 60-minutes pulse labelling, we observed a robust inhibition of mitochondrial protein synthesis for all of the snMRP^KO^ cell lines except mS37^KO^ (Figure 4A). However, an extended labelling reaction for 4 hours did reveal low levels of mitochondrial protein synthesis in mS29^KO^, mS33^KO^ and mS35^KO^ cell lines (Figure S6).

**Figure 4.**
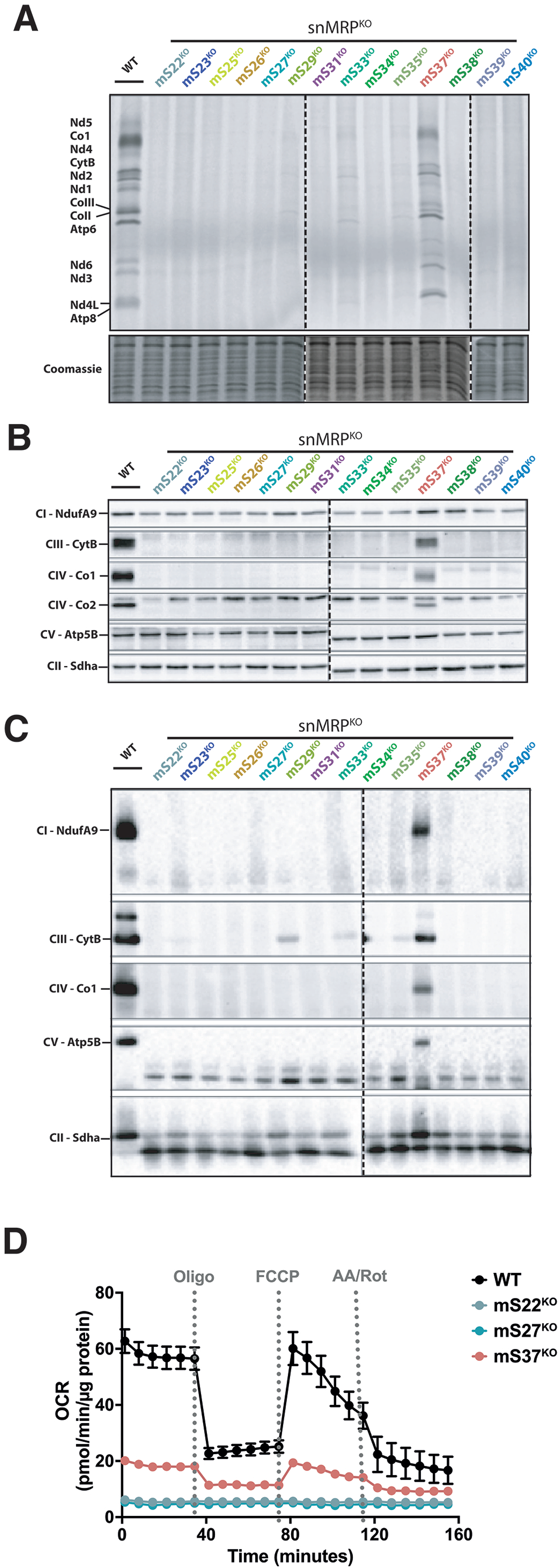
Effect of snMRP knockout on mitochondrial protein synthesis and steady-state of OXPHOS complexes and subunits. **(A)** ^35^S-metabolic labelling to assess mitochondrial protein synthesis in snMRP^KO^ lines. **(B)** Immunoblot of steady-state mitochondrial OXPHOS complex subunits. **(C)** Immunoblot of native OXPHOS complexes. Dotted line delineates separate gel where controls are excluded for readability. **(D)** Seahorse High-resolution respirometry in selected snMRP knockout lines.

Next, we investigated the expression and assembly of oxidative phosphorylation complexes, using SDS and Blue-Native PAGE analyses (Figure 4B, 4C) in all our snMRP^KO^ cell lines. These analyses revealed undetectable levels of the assembled complexes in most snMRP^KO^ cell lines except mS37^KO^. Although in mS29^KO^, mS33^KO^ and mS35^KO^ cell lines some residual level of assembled complex III could be detected. High-resolution respirometry revealed mS37^KO^ to have a residual oxidative phosphorylation capacity (Figure 4D).

### The stability of mS37 is dependent on oxidizable cysteine residues for regulating mitochondrial protein synthesis

Of all the snMRP^KO^ cell lines that we generated, only mS37^KO^ retained substantial mitochondrial protein synthesis and OXPHOS complexes, and yet mS37 stability was reduced in all of these cell lines (Figure 1B). This result was surprising. In contrast, the abundance of the other mtSSU proteins was determined by the presence or absence of neighbouring interacting proteins. To understand the role of mS37 in mammalian translation further, we generated two knockout cell lines of snMRPs in the mtLSU to test if the protein stability correlated with assembly defects and/or overall defects in mitochondrial protein synthesis (Figure 5A). Knockouts for mL44^KO^, a known disease associated mtLSU protein (Carroll et al., 2013), and mL46^KO^ generated a defect in mitochondrial protein synthesis (Figure 5B) and a reduction in the steady-state abundance of mitochondrially–encoded OXPHOS complexes (Figure 5C). Immunoblotting, isokinetic sucrose gradients, and immunofluorescence staining of mS37 of these snMRP^KO^ cell lines revealed that whilst the mtSSU remains intact in the mtLSU knockouts, the stability of mS37 is impaired (Figure 5D-F). A recent report using proximity labelling with BioID suggests that mS37 could be dual localised within the cell (Antonicka et al., 2020). However, immunofluorescence experiments with mS37 demonstrate only a mitochondrial localisation (Figure 5F). Together, these findings show that mS37 stability is compromised in assembly defects of both the mtSSU and mtLSU but on its own is not essential for assembly.

**Figure 5.**
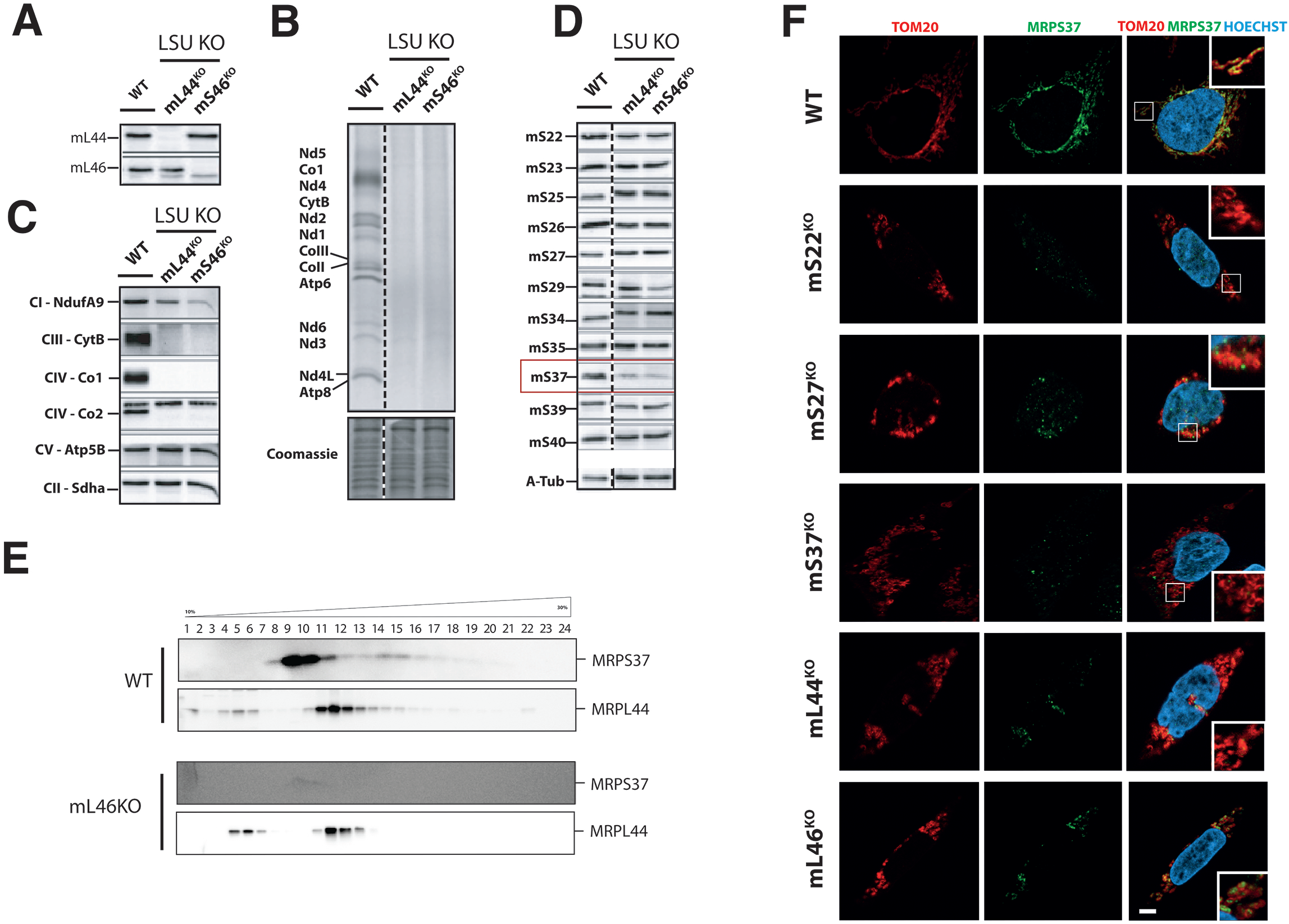
Effect of mtLSU knockouts on mS37 stability. **(A)** Immunoblot confirming CRISPR-mediated mL44^KO^ and mL46^KO^. **(B)** ^35^S-metabolic labelling to assess mitochondrial protein synthesis in mL44^KO^ and mL46^KO.^ **(C)** Immunoblot for steady-state of OXPHOS complex subunits in mL44^KO^ and mL46^KO^. **(D)** Immunoblot confirming impaired mS37 stability in mL44^KO^ and mL46^KO^. **(E)** Isokinetic sucrose gradients of mitoribosome assembly in mL46^KO^. **(F)** Immunofluorescence to assess mitochondrial structure and mS37 localisation using antibodies against TOM20 and mS37. DNA stained with Hoechst label. Right panel shows overlay of TOM20, mS37 and Hoechst with inset showing magnified area in box. Scale bar = 2 μm.

One interpretation of the mS37 phenotypes is that the protein has a regulatory function in mitochondrial protein synthesis. mS37 has a coiled-coil-helix-coiled-coil-helix domain (CHCHD) containing conserved twin CX_9_C motifs (Habich et al., 2019) (Figure 6A-B). Mitochondrial import of the CHCHD family members is regulated by the MIA40 receptor, which forms disulfide bonds at the cysteine residues to stabilise the nascent protein during import (Longen et al., 2014). This interaction is essential for mitochondrial protein import with factors that lack classical mitochondrial targeting sequences (Habich et al., 2019). Defects in this redox mediated regulation, leads to protein import failures (Habich et al., 2019). (Finger and Riemer, 2020; Longen et al., 2014). To test whether the CX_9_C motif in mS37 is necessary for protein stability, we mutated all four cysteines to serine within the CX_9_C motif (Figure 6C) and expressed a FLAG-tagged version (mS37 C1234S) in the mS37^KO^ cell line. Whilst wild-type mS37 was stable the mutant mS37 C1234S was detectable only at very low levels, suggesting the cysteine residues were necessary for protein stability (Figure 6D).

**Figure 6.**
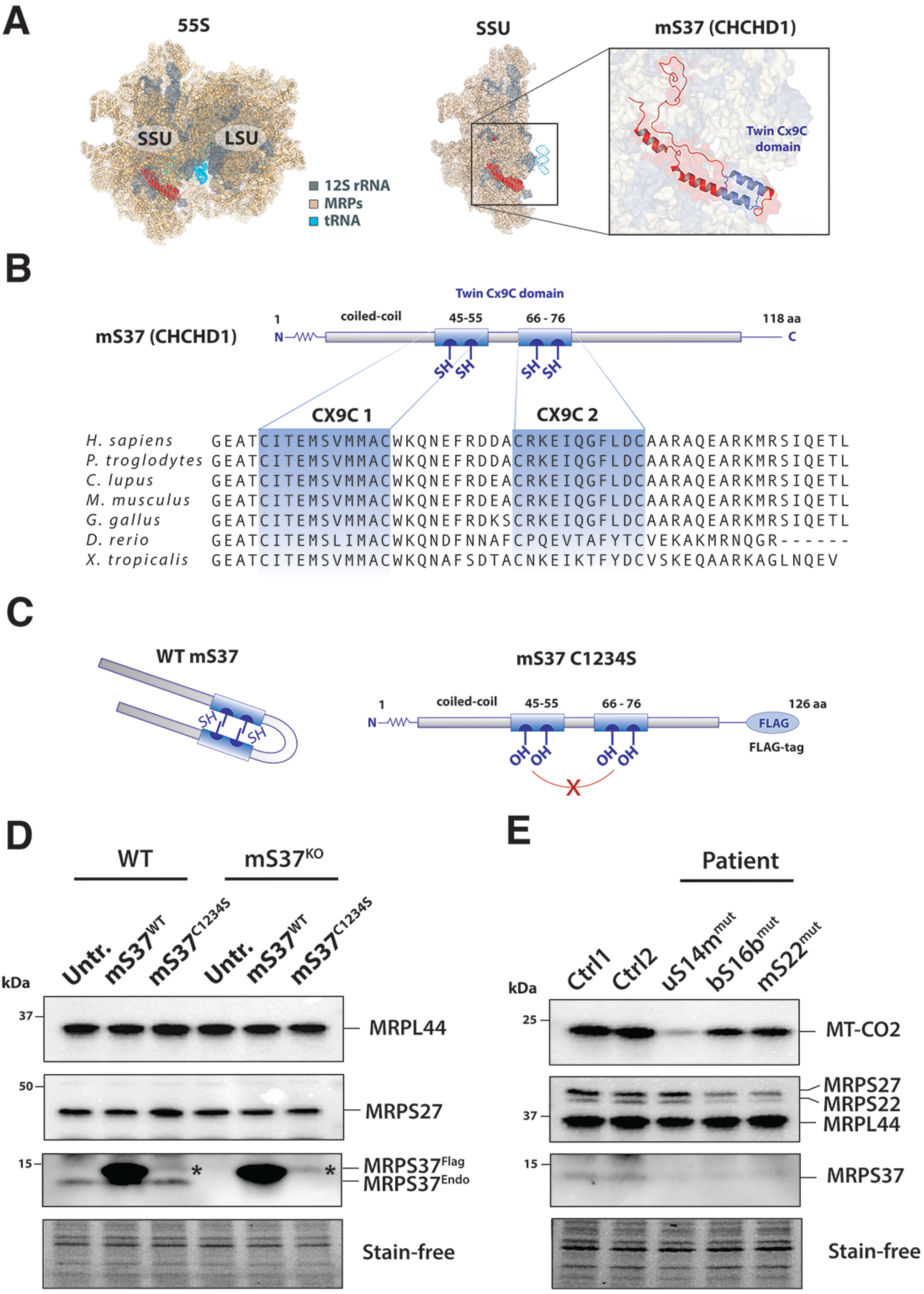
Stability of mS37 is regulated by CX_9_C motif. **(A)** Location of mS37 within the 55S mitoribosome. **(B)** Conservation of the CHCHD4-oxidisable twin CX_9_C motifs in mS37 and (**C**) subsequent folding. **(C)** Schematic representation of 4 mutated cysteine residues into serine (C1234S). **(D)** Immunoblot of FLAG-tagged wildtype and C1234S mS37. **(E)** Immunoblot of mS37 protein level in patient-derived fibroblasts.

We hypothesised that stability mS37 would be impaired in diseases associated with variants in mtSSU proteins. Therefore, we examined the stability of mS37 in patient-derived fibroblasts with mitochondrial ribosomal pathogenic variants in the mtSSU (uS14m, bS16b and mS22) (Jackson et al., 2019; Miller et al., 2004; Saada et al., 2007). In all cases, mS37 levels were reduced compared to controls (Figure 6E). Together our results suggest mS37 is acutely sensitive to defects in mitochondrial protein synthesis and that its stability is regulated via disulfide bonds.

## Discussion

In this study we investigated the importance of the human snMRPs of the mtSSU to mitochondrial ribosome assembly. We show that most mitochondrial-specific snMRPs are essential for mtSSU assembly and mitochondrial protein synthesis, with the exception of mS37 (MRPS37/CHCHD1). This highlights a unique mtSSU protein with a highly specialised role in mitochondrial protein synthesis. Our experiments support that mS37 could act as maturation control of the ribosome prior to the initiation of translation. Recently, cryo-electron microscopic structures determined translation pre-initiation steps of the mtSSU and revealed that mS37 and mtIF3 interaction provides a conformation favourable for mtIF2 accommodation (Khawaja et al., 2020), and mS37 links the final steps of mtSSU assembly and mitochondrial translation initiation (Itoh et al., 2022). Moreover, we found oxidisable cysteine residues in a twin CX_9_C motif in mS37 to be critical for its stability, suggesting redox-regulated mS37 stability as a mechanism for regulating mitochondrial translation initiation.

In addition to its proposed function in the mtSSU, a proximity-labelling approach showed mS37 to have strong interactions with nucleolar and nuclear envelope proteins, suggesting potential multi-compartment targeting of mS37 (Antonicka et al., 2020). Furthermore, evidence for dual-localisation of human mS37 is also supported by a study showing wild-type protein and a C-terminally truncated version with nuclear localisation (Westerman et al., 2004). We could not confirm nuclear localisation of mS37 with immunofluorescence, and therefore the function of nuclear localised mS37 proteins remains to be resolved.

By assessing mtSSU protein abundance, we observed clusters loosely fitting previously reported assembly modules determined by pulse-labelling proteomic methods (Bogenhagen et al., 2018). This confirms that individual gene knockouts can be leveraged to uncover the mitoribosome assembly pathway (Zeng et al., 2018). Our analyses revealed the identities of closely interacting proteins which likely form the modular units, refining the predictions made based on data from pulse-labelling methods (Bogenhagen et al., 2018).

Although all MRPs seem stably expressed through tissue development, they show differential expression in certain cancers and disease presentations (Cheong et al., 2020; Jackson et al., 2019; Kim et al., 2017; Mays et al., 2019). This suggests the potential of heterogenous mitoribosome composition across cell types. Indeed, mutations in mitoribosomal proteins lead to variable disease phenotypes (Table S1). To date, snMRP proteins associated with human disease are mS22, mS23, mS25, mS34 and mS39 (Saada et al., 2007; Bugiardini et al., 2019; Lake et al., 2017; Borna et al., 2019). All reported patients have mitoribosome instability. Although the pathomechanistic consequence is combined OXPHOS deficiency, pathogenic variants in snMRPs lead to distinct phenotypic manifestations. This includes, brain abnormalities and hypertrophic cardiomyopathy for mS22 (Saada et al., 2007) hepatic disease for mS23 (Kohda et al., 2016), encephalomyopathy for mS25 (Bugiardini et al., 2019), Leigh syndrome for mS34 (Lake et al., 2017) and mS39 (Borna et al., 2019).

In summary, we highlight mS37 as a unique protein of the mtSSU in regulating mitochondrial protein synthesis and demonstrate the essential nature of snMRPs for mtSSU assembly and OXPHOS. Furthermore, we identify the composition of proteins that form modular units in the mtSSU assembly pathway.

## Author contribution

T.H., C.B.J., C.J.C. conceived and designed the study. T.H., G.M., R.A., M.P., K.B., T.N., I.C., D.B., C.B.J., C.J.C. conducted experiments. T.H., G.M., R.A., C.B.J., T.N., E.P., analysed the data. B.J.B. provided critical reagents and editing of the manuscript. T.H., C.B.J. and C.J.C. wrote the manuscript, approved by all authors.

## Acknowledgments and Funding

The authors wish to thank Tarja Grundström for technical help. Following funding resources are acknowledged: The Lily Foundation [K.B] [C.J.C], Academy of Finland [C.B.J], the Magnus Ehrnroot Foundation [C.B.J], the ILS Doctoral Programme [R.A.], Säätiöiden post doc-pooli, Paulon säätiö [T.H.], Maud Kuistila Memorial Foundation [T.H.], Emil Aaltosen säätiö [T.H.], EMBO Long-Term Fellowship ALT295-2019 [D.B.] BJB was supported by the Academy of Finland (307431 and 314706), the Sigrid Juselius Foundation Senior Investigator Award, and donations from Hereditary Neuropathy Foundation and Lindsey Flynt. Mass spectrometry-based proteomic analyses were performed by the Proteomics Core Facility, Department of Immunology, University of Oslo/Oslo University Hospital, which is supported by the Core Facilities program of the South-Eastern Norway Regional Health Authority. This core facility is also a member of the National Network of Advanced Proteomics Infrastructure (NAPI), which is funded by the Research Council of Norway INFRASTRUKTUR-program (project number: 295910). We acknowledge the use of the Image Resource Facility, St George’s, University of London.

## Materials and Methods

### Design and generation of gRNA-oligos

20 base pair CRISPR-RNA (crRNA) sequences were designed with Benchling (https://www.benchling.com). Two gRNAs per gene were used to preferentially excise the start codon site. The crRNAs were selected according to their off- and on-target score predictions. Testing of the efficiency of crRNAs was performed as described before by the use of a three tailed-template PCR (Balboa et al., 2015). Briefly, the 20 bp crRNAs were combined from the 5’ end to an RNA oligo containing 19 bp matching to a tailed U6 promoter plus an extra G nucleotide (for proper RNA transcription), and from the 3’ end to an RNA oligo containing 19bp matching to a tracRNA and tailed terminator as follows (5’-GTGGAAAGGACGAAACACCgNNNNNNNNNNNNNNNNNNNNGTTTTAGAGC TAGAAATAG-3’). The 59 bp gRNA-oligos were ordered from Sigma. The gRNA-oligos used are listed in Supplementary Table S2. Next, U6 promoter containing plasmid pSpCas9(BB)-2A-GFP (PX458) (Addgene, #48138) was digested with the BbsI and PvuI and run on 1% agarose gel. The ∼1.5 kb band was excised from the gel and column-purified. The purified U6 promoter was used as a template in a PCR using Phusion enzyme (ThermoScientific, F530S). The resulting PCR product was column-purified and digested with BbsI overnight to get rid of any possible remaining plasmid. The PCR product was separated on a 1% agarose gel and the 249 bp fragment excised and column-purified.

Using a tailed-PCR-assay, gRNA oligonucleotides were amplified with a U6 promoter and mTERM fragment as previously established (Balboa et al., 2015). Tailed U6 was produced by using the previously digested U6 promoter as a template with primers 5ptailedU6promFw and U6promRv and tailed terminator was produced by using Long reverse oligo TermRv80bp as a template with primers Term80bp and 3ptailedterm80bpRv in a PCR (primers listed in Supplementary table S3). Resulting products were run on 1 % agarose and the 266 bp PCR product for tailed U6 and 103 bp PCR product for tailed terminator were excised and column-purified. Transcriptional gRNA units were prepared by PCR from gRNA-oligos and tailed U6 and terminators in a PCR.

### Testing and transfecting cells with gRNAs

Wild-type HEK293 cells were seeded on 24-well plates for ∼ 40% confluency the day before transfections. On the day of transfection, media was changed, and cells were transfected with 500 ng of CAG-Cas9-T2A-EGFP-ires-puro-plasmid (Addgene, #78311) and with 250 ng of gRNAs altogether using PEI transfection reagent with addition of 0.15 M NaCl. The media was changed one day after transfection. Cells were transfected altogether for two days. Transfected cells were FACS-sorted using the co-transfected GFP-CAS9 reporter and knockout clones expanded from GFP-positive FACS-sorted single cells.

### DNA extraction and PCR confirmation

DNA was extracted from HEK-293 cells using DirectPCR® DNA Lysis Reagent (VWR, 732-3255) according to the manufacturer’s instructions. PCR confirmation of the gRNA transfected cells and clonal cell lines were performed in a standard PCR with Sanger sequencing (primers listed in Supplementary table S4).

### mS37 plasmid transfection

Plasmids expressing wild-type or mutated mS37 cDNAs were purchased from OriGene. HEK293 cells (wild-type or mS37 knock-out) were transfected in 6-well cell culture dish at 70% confluency with 2 μg plasmid using Lipofectamine 2000 (ThermoFisher scientific, Cat#11668027). Cells were collected after 72h, lysed with RIPA buffer, and analysed by immunoblotting (see section “immunoblotting”).

### Immunoblotting

Cells were lysed in 1xPBS containing Pierce™ EDTA-free Protease Inhibitor Mini Tablets (ThermoFisher Scientific, A32955), 1% N-Dodecyl-b-D-Maltoside (Amresco, J424,), 1% Phenylmethanesulfonyl fluoride (PMSF) (Sigma, 93482) on ice for 30 min with subsequent centrifugation at 14000g at 4°C for 25 min. 20 μg of the resulting protein lysates were mixed with 1 x Laemmli loading dye containing 5% 2-Mercaptoethanol (Sigma, M3148), separated on 10% polyacrylamide gel and semi-dry blotted to nitrocellulose membrane. The membranes were blocked in TBS-T with 5% milk powder for 1h, incubated in primary antibody overnight at 4°C and detected with the respective HRP-secondaries using chemiluminescence. Antibodies are listed in Supplementary table S5.

### Blue-Native acrylamide electrophoresis

10 cm diameter plates of cells were washed with PBS and scraped into 450 μl of ice-cold PBS. Protein concentrations were determined using Bradford assay and the cells were pelleted by centrifugation in 10,000g for 10 minutes at 4°C. Cell pellets were resuspended in MB2 buffer, for final protein concentration of 3 μg/μl (for buffers see (Schägger and Jagow, 1991)). n-Dodecyl-β-Maltoside (DDM) was added for a final concentration of 1%, samples incubated on ice for 30 min and centrifuged at 20,000g for 20 minutes at 4°C, after which the supernatant was transferred to a new tube, protein concentration determined, and appropriate loading dye added. 15 μg of protein was separated on NativePAGE 3-12% Bis/Tris gel (ThermoFisher, BN1003BOX) and semi-dry blotted to a nitrocellulose membrane and detected as described above.

### Translation assays

The cells were grown on 60 mm diameter dishes and labelling was performed at 80 % cell confluency. The cells were washed once with 1xPBS and incubated in pre-warmed labelling medium (DMEM without methionine and cysteine (Sigma, D0422), 1xglutamax (Gibco, 35050-038), 10% dialysed serum (Thermo Fisher Scientific, A3382001) and 50 μg/ml uridine (Sigma, U3003) for 25 min. 100 μg/ml of anisomycin (Sigma, A9789) was added to each plate 5 minutes prior to addition of labeling mix to inhibit cytoplasmic translation. After that 200μCi/ml of 35S-methionine/cysteine (PerkinElmer, EasyTag EXPRESS ^35^S Protein Labeling Mix NEG072014MC) was added to each plate and the plates were incubate for 30 minutes. Cells were pulse labelled for 1 or 4 hours after which the cells were washed twice with 1xPBS and scraped into 1 ml of ice-cold 1xPBS. The cells were collected by centrifugation in 14‵000g for 10 minutes at 4°C. The cells were resuspended in ice-cold 1xPBS depending on the pellet size and protein concentration was measured using Bradford. 30 μg of protein was span down by 14,000g, for 20 minutes at 4°C. The protein pellets were resuspended in 10 μl of H_2_O and 0.3 μl of Benzonase (≥250 units/μl, Sigma, E1014)/sample and the samples were incubated in RT for 5 min. After that 20 μl of 2xloading buffer (186 mM Tris-HCl, pH=6.7-6.8, 15% glycerol, 2% SDS, 0.5 mg/ml bromophenol blue, 6% β-mercaptoethanol) was added and the samples were equilibrated in RT for 1 hour. The samples were run into 12-20% gradient polyacrylamide gel overnight. Next day the gel was rinsed with milliQ-water and dried at 60°C in a gel drier for 75 minutes. The gel was exposed to a phosphor screen and the screen was imaged using Typhoon 9400 scanner.

### Sucrose gradients

Sedimentation of mitoribosomes was studied by sucrose gradients. Cells were lysed with 1% DDM lysis buffer (50 mM Tris, pH 7.2, 10 mM Mg(CH_3_COO)_2_, 40 mM NH_4_Cl, 100 mM KCl, 1% DDM, 1 mM PMSF, 1,24 mM chloramphenicol, and 1 mM ATP) by incubating the samples for 20 minutes on ice followed by centrifugation at 20‵000xg for 20 minutes at 4°C. Protein concentration in the supernatant was determined by Bradford assay and 2 mg of protein of each sample was loaded on top of a 16 ml linear 10-30% sucrose gradient (50 mM Tris, pH 7.2, 10 mM Mg(CH_3_COO)_2_, 40 mM NH_4_Cl, 100 mM KCl, 1 mM PMSF, and 1 mM ATP). The tubes were centrifuged for 15 hours at 4°C and 74‵400g. The gradients were divided into 24 fractions of equal volume from top to bottom and the fractions were TCA precipitated. Samples were separated in 12–20% gradient Tris-Glycine SDS-PAGE gels, immunoblotted and detected as described above. Gels were rehydrated in water and Coomassie stained to confirm loading.

### High-resolution respirometry

Mitochondrial respiration was measured using an XFe96 Extracellular Flux Analyzer (Agilent). Cells were cultured in assay-specific 96-well culture plates optimized for cell density (8,000 cells/well). One hour prior to measurement, culture medium was replaced with XF DMEM assay medium (Agilent, 103680-100), supplemented with 10 mM glucose (Sigma, G8270,), 1 mM sodium pyruvate (ThermoFisher Scientific, 11360070,), and 2 mM L-glutamine (ThermoFisher Scientific, 25030123) after which cells were incubated in a non-CO_2_ 37 °C incubator. The assay consisted of assessment of basal respiration, after which 1.5 μM oligomycin (Sigma, O4876) was added to measure leak respiration, 1.125 μM carbonylcyanide-4-(trifluoromethoxy)-phenylhydrazone (FCCP (Sigma, C2920)) to measure maximal uncoupled respiration) and a mixture of 1.0 μM antimycin A (Sigma, A8674) and rotenone (Sigma, R8875) to measure residual respiration.

### Immunofluorescence experiments and analysis

Cells were cultured on coverslips and fixed with 4% paraformaldehyde for 10 minutes. The fixed samples were then washed with PBS for 10 minutes. The cells were permeabilised with 0.1% TritonX-100 for 30 minutes, blocked in 10% horse serum and washed with PBS for 5 minutes. The samples were incubated overnight at 4°C with the following primary antibodies: mouse anti-TOM20 (SantaCruz, sc-11415, 1:400), rabbit anti-MRPS37/mS37 (Proteintech, 11728-1-AP, 1: 400). Next, the cells were washed with PBS for 5 minutes and incubated for 30 minutes with the following secondary antibodies: goat anti-rabbit AlexaFluor 488 (Invitrogen Antibodies, A-21207, 1: 500) and goat anti-mouse AlexaFluor 594 (Invitrogen Antibodies, A-21206, 1: 500). The cells were washed twice with 1xPBS for 5 minutes, stained with 1xPBS containing Hoechst for 5 minutes, washed twice with 1xPBS for 15 minutes and observed with the Zeiss Axio Observer Z1 inverted phase contrast fluorescence microscope with the peak emission wavelengths of 618 nm (red), 517 nm (green) and 465 nm (blue). Mitochondrial fragmentation and co-localization analysis were performed with a minimum of 100 cells per group.

### Label-free quantitative proteomics

Proteomic analysis was performed in quadruplicates for each cell line. Briefly, each sample was lysed with RIPA-buffer, proteins were precipitated with acetone and trypsin-digested using sequencing grade-modified trypsin (Promega). The resulting peptide mixture was purified by STAGE-TIP method using a C18 resin disk (3M Empore) before the samples were analysed by nanoLC-MS/MS using QExactive HF (Thermo) coupled to nEASY-LC (Thermo).

MS raw files were submitted to MaxQuant software v.1.6.1.0 for protein identification and label-free quantification. Carbamidomethyl (C) was set as a fixed modification and acetyl (protein N-term), carbamyl (N-term) and oxidation (M) were set as variable modifications. First search peptide tolerance of 20 ppm and main search error 4.5 ppm were used. Trypsin without proline restriction enzyme option was used, with two allowed miscleavages. The minimal unique + razor peptides number was set to 1, and the allowed FDR was 0.01 (1%) for peptide and protein identification. Label-free quantitation was employed with default settings. UniProt database with ‘Human’ entries (2018) was used for the database searches. Known contaminants as provided by MaxQuant and identified in samples were excluded from further analysis. Perseus software v.1.6.1.3 was used for statistical analysis of the label-free quantification data. The mass spectrometry proteomics data have been deposited to the ProteomeXchange Consortium via the PRIDE partner repository with the dataset identifier PXD034224.

### Statistics and software

All graphical representations were performed in Adobe Illustrator with statistic calculations performed in Prism. Structural representations were created using Pymol v.2.4.1.

## References

Aibara, S., Singh, V., Modelska, A., and Amunts, A. (2020). Structural basis of mitochondrial translation. Elife 9, 531. https://doi.org/10.7554/elife.58362.

Amunts, A., Brown, A., Toots, J., Scheres, S.H.W., and Ramakrishnan, V. (2015). Ribosome. The structure of the human mitochondrial ribosome. Science 348, 95–98. https://doi.org/10.1126/science.aaa1193.

Antonicka, H., Lin, Z.-Y., Janer, A., Aaltonen, M.J., Weraarpachai, W., Gingras, A.-C., and Shoubridge, E.A. (2020). A High-Density Human Mitochondrial Proximity Interaction Network. Cell Metabolism 32, 479–497.e9. https://doi.org/10.1016/j.cmet.2020.07.017.

Balboa, D., Weltner, J., Eurola, S., Trokovic, R., Wartiovaara, K., and Otonkoski, T. (2015). Conditionally Stabilized dCas9 Activator for Controlling Gene Expression in Human Cell Reprogramming and Differentiation. Stem Cell Reports 5, 448–459. https://doi.org/10.1016/j.stemcr.2015.08.001.

Ban, N., Beckmann, R., Cate, J.H.D., Dinman, J.D., Dragon, F., Ellis, S.R., Lafontaine, D.L.J., Lindahl, L., Liljas, A., Lipton, J.M., et al. (2014). A new system for naming ribosomal proteins. Current Opinion in Structural Biology 24, 165–169. https://doi.org/10.1016/j.sbi.2014.01.002.

Bogenhagen, D.F., Ostermeyer-Fay, A.G., Haley, J.D., and Garcia-Diaz, M. (2018). Kinetics and Mechanism of Mammalian Mitochondrial Ribosome Assembly. Cell Rep 22, 1935–1944. https://doi.org/10.1016/j.celrep.2018.01.066.

Borna, N.N., Kishita, Y., Kohda, M., Lim, S.C., Shimura, M., Wu, Y., Mogushi, K., Yatsuka, Y., Harashima, H., Hisatomi, Y., et al. (2019). Mitochondrial ribosomal protein PTCD3 mutations cause oxidative phosphorylation defects with Leigh syndrome. Neurogenetics 1–17. https://doi.org/10.1007/s10048-018-0561-9.

Brown, A., Amunts, A., Bai, X., Sugimoto, Y., Edwards, P.C., Murshudov, G., Scheres, S.H.W., and Ramakrishnan, V. (2014). Structure of the large ribosomal subunit from human mitochondria. Science 346, 718–722. https://doi.org/10.1126/science.1258026.

Bugiardini, E., Mitchell, A.L., Rosa, I.D., Horning-Do, H.-T., and Spinazzola, A. (2019). MRPS25 mutations impair mitochondrial translation and cause encephalomyopathy. Hum. Mol. Genet. 1–10. https://doi.org/10.1093/hmg/ddz093ddz093.

Carroll, C.J., Isohanni, P., Pöyhönen, R., Euro, L., Richter, U., Brilhante, V., Götz, A., Lahtinen, T., Paetau, A., Pihko, H., et al. (2013). Whole-exome sequencing identifies a mutation in the mitochondrial ribosome protein MRPL44 to underlie mitochondrial infantile cardiomyopathy. J. Med. Genet. 50, 151–159. https://doi.org/10.1136/jmedgenet-2012-101375.

Cheong, A., Lingutla, R., and Mager, J. (2020). Expression analysis of mammalian mitochondrial ribosomal protein genes. Gene Expression Patterns 38, 119147. https://doi.org/10.1016/j.gep.2020.119147.

Desai, N., Brown, A., Amunts, A., and Ramakrishnan, V. (2017). The structure of the yest mitochondria ribosome. Science 355, 528–531. https://doi.org/10.1126/science.aal2415.

Desai, N., Yang, H., Chandrasekaran, V., Kazi, R., Minczuk, M., and Ramakrishnan, V. (2020). Elongational stalling activates mitoribosome-associated quality control. Science 370, 1105–1110. https://doi.org/10.1126/science.abc7782.

Ferrari, A., Del’Olio, S., and Barrientos, A. (2020). The Diseased Mitoribosome. FEBS Letters 233, 657–37. https://doi.org/10.1002/1873-3468.14024.

Finger, Y., and Riemer, J. (2020). Protein import by the mitochondrial disulfide relay in higher eukaryotes. Biol. Chem. 401, 749–763. https://doi.org/10.1515/hsz-2020-0108.

Galmiche, L., Serre, V., Beinat, M., Assouline, Z., Lebre, A.-S., Chrétien, D., Nietschke, P., Benes, V., Boddaert, N., Sidi, D., et al. (2011). Exome sequencing identifies MRPL3 mutation in mitochondrial cardiomyopathy. Hum. Mutat. 32, 1225–1231. https://doi.org/10.1002/humu.21562.

Gardeitchik, T., Mohamed, M., Ruzzenente, B., Karall, D., Guerrero-Castillo, S., Dalloyaux, D., Brand, M. van den, Kraaij, S. van, Asbeck, E. van, Assouline, Z., et al. (2018). Bi-allelic Mutations in the Mitochondrial Ribosomal Protein MRPS2 Cause Sensorineural Hearing Loss, Hypoglycemia, and Multiple OXPHOS Complex Deficiencies. Am. J. Hum. Genet. 102, 685–695. https://doi.org/10.1016/j.ajhg.2018.02.012.

Greber, B.J., Bieri, P., Leibundgut, M., Leitner, A., Aebersold, R., Boehringer, D., and Ban, N. (2015). Ribosome. The complete structure of the 55S mammalian mitochondrial ribosome. Science 348, 303–308. https://doi.org/10.1126/science.aaa3872.

Habich, M., Salscheider, S.L., and Riemer, J. (2019). Cysteine residues in mitochondrial intermembrane space proteins. Br J Pharmacol 176, 514–531. https://doi.org/10.1111/bph.14480.

Hart, T., Chandrashekhar, M., Aregger, M., Steinhart, Z., Brown, K.R., MacLeod, G., Mis, M., Zimmermann, M., Fradet-Turcotte, A., Sun, S., et al. (2015). High-Resolution CRISPR Screens Reveal Fitness Genes and Genotype-Specific Cancer Liabilities. Cell 163. https://doi.org/10.1016/j.cell.2015.11.015.

Hilander, T., Jackson, C.B., Robciuc, M., Bashir, T., and Zhao, H. (2021). The Roles of Assembly Factors in Mammalian Mitoribosome Biogenesis. Mitochondrion 60, 70–84. https://doi.org/10.1016/j.mito.2021.07.008.

Itoh, Y., Andréll, J., Choi, A., Richter, U., Maiti, P., Best, R.B., Barrientos, A., Battersby, B.J., and Amunts, A. (2021). Mechanism of membrane-tethered mitochondrial protein synthesis. Science 371, 846–849. https://doi.org/10.1126/science.abe0763.

Itoh, Y., Khawaja, A., Laptev, I., Cipullo, M., Atanassov, I., Sergiev, P., Rorbach, J., and Amunts, A. (2022). Mechanism of mitoribosomal small subunit biogenesis and preinitiation. Nature 1–6. https://doi.org/10.1038/s41586-022-04795-x.

Jackson, C.B., Gallati, S., and Schaller, A. (2012). qPCR-based mitochondrial DNA quantification: Influence of template DNA fragmentation on accuracy. BIOCHEMICAL AND BIOPHYSICAL RESEARCH COMMUNICATIONS 1–7. https://doi.org/10.1016/j.bbrc.2012.05.121.

Jackson, C.B., Zbinden, C., Gallati, S., and Schaller, A. (2014). Heterologous expression from the human D-Loop in organello. Mitochondrion 17, 67–75. https://doi.org/10.1016/j.mito.2014.05.011.

Jackson, C.B., Huemer, M., Bolognini, R., Martin, F., Szinnai, G., Donner, B.C., Richter, U., Battersby, B.J., Nuoffer, J.-M., Suomalainen, A., et al. (2019). A variant in MRPS14 (uS14m) causes perinatal hypertrophic cardiomyopathy with neonatal lactic acidosis, growth retardation, dysmorphic features and neurological involvement. Hum. Mol. Genet. 28, 639–649. https://doi.org/10.1093/hmg/ddy374.

Khawaja, A., Itoh, Y., Remes, C., Spahr, H., Yukhnovets, O., Höfig, H., Amunts, A., and Rorbach, J. (2020). Distinct pre-initiation steps in human mitochondrial translation. Nat Commun 1–10. https://doi.org/10.1038/s41467-020-16503-2.

Kim, H.-J., Maiti, P., and Barrientos, A. (2017). Mitochondrial ribosomes in cancer. Seminars in Cancer Biology 47, 67–81. https://doi.org/10.1016/j.semcancer.2017.04.004.

Koc, E.C.K. and H. (2013). Identification and characterization of CHCHD1, AURKAIP1, and CRIF1 as new members of the mammalian mitochondrial ribosome. 1–15. https://doi.org/10.3389/fphys.2013.00183/abstract.

Kohda, M., Tokuzawa, Y., Kishita, Y., Nyuzuki, H., Moriyama, Y., Mizuno, Y., Hirata, T., Yatsuka, Y., Yamashita-Sugahara, Y., Nakachi, Y., et al. (2016). A Comprehensive Genomic Analysis Reveals the Genetic Landscape of Mitochondrial Respiratory Chain Complex Deficiencies. PLoS Genet. 12, e1005679–31. https://doi.org/10.1371/journal.pgen.1005679.

Koripella, R.K., Sharma, M.R., Bhargava, K., Datta, P.P., Kaushal, P.S., Keshavan, P., Spremulli, L.L., Banavali, N.K., and Agrawal, R.K. (2020). Structures of the human mitochondrial ribosome bound to EF-G1 reveal distinct features of mitochondrial translation elongation. Nat Commun 1–12. https://doi.org/10.1038/s41467-020-17715-2.

Kummer, E., and Ban, N. (2021). Mechanisms and regulation of protein synthesis in mitochondria. Nat. Rev. Mol. Cell Biol. 1–19. https://doi.org/10.1038/s41580-021-00332-2.

Lake, N.J., Webb, B.D., Stroud, D.A., Richman, T.R., Ruzzenente, B., Compton, A.G., Mountford, H.S., Pulman, J., Zangarelli, C., Rio, M., et al. (2017). Biallelic Mutations in MRPS34 Lead to Instability of the Small Mitoribosomal Subunit and Leigh Syndrome. The American Journal of Human Genetics 101, 239–254. https://doi.org/10.1016/j.ajhg.2017.07.005.

Longen, S., Woellhaf, M.W., Petrungaro, C., Riemer, J., and Herrmann, J.M. (2014). The Disulfide Relay of the Intermembrane Space Oxidizes the Ribosomal Subunit Mrp10 on Its Transit into the Mitochondrial Matrix. Dev. Cell 28, 30–42. https://doi.org/10.1016/j.devcel.2013.11.007.

Mays, J.-N., Camacho-Villasana, Y., Garcia-Villegas, R., Perez-Martinez, X., Barrientos, A., and Fontanesi, F. (2019). The mitoribosome-specific protein mS38 is preferentially required for synthesis of cytochrome c oxidase subunits. Nucleic Acids Research 47, 5746–5760. https://doi.org/10.1093/nar/gkz266.

Menezes, M.J., Guo, Y., Zhang, J., Riley, L.G., Cooper, S.T., Thorburn, D.R., Li, J., Dong, D., Li, Z., Glessner, J., et al. (2015). Mutation in mitochondrial ribosomal protein S7 (MRPS7) causes congenital sensorineural deafness, progressive hepatic and renal failure and lactic acidemia. Hum. Mol. Genet. 24, 2297–2307. https://doi.org/10.1093/hmg/ddu747.

Miller, C., Saada, A., Shaul, N., Shabtai, N., Ben-Shalom, E., Shaag, A., Hershkovitz, E., and Elpeleg, O. (2004). Defective mitochondrial translation caused by a ribosomal protein (MRPS16) mutation. Ann. Neurol. 56, 734–738. https://doi.org/10.1002/ana.20282.

Nottia, M.D., Marchese, M., Verrigni, D., Mutti, C.D., Torraco, A., Oliva, R., Fernandez-Vizarra, E., Morani, F., Trani, G., Rizza, T., et al. (2020). A homozygous MRPL24 mutation causes a complex movement disorder and affects the mitoribosome assembly. Neurobiol Dis 141, 104880. https://doi.org/10.1016/j.nbd.2020.104880.

Pulman, J., Ruzzenente, B., Bianchi, L., Rio, M., Boddaert, N., Munnich, A., Rötig, A., and Metodiev, M.D. (2018). Mutations in the MRPS28gene encoding the small mitoribosomal subunit protein bS1m in a patient with intrauterine growth retardation, craniofacial dysmorphism and multisystemic involvement. Hum. Mol. Genet. 515, 283–18. https://doi.org/10.1093/hmg/ddy441.

Rackham, O., and Filipovska, A. (2014). Supernumerary proteins of mitochondrial ribosomes. Biochimica Et Biophysica Acta Bba - Gen Subj 1840, 1227–1232. https://doi.org/10.1016/j.bbagen.2013.08.010.

Saada, A., Shaag, A., Arnon, S., Dolfin, T., Miller, C., Fuchs-Telem, D., Lombès, A., and Elpeleg, O. (2007). Antenatal mitochondrial disease caused by mitochondrial ribosomal protein (MRPS22) mutation. J. Med. Genet. 44, 784–786. https://doi.org/10.1136/jmg.2007.053116.

Saurer, M., Ramrath, D.J.F., Niemann, M., Calderaro, S., Prange, C., Mattei, S., Scaiola, A., Leitner, A., Bieri, P., Horn, E.K., et al. (2019). Mitoribosomal small subunit biogenesis in trypanosomes involves an extensive assembly machinery. Science 365, 1144–1149. https://doi.org/10.1126/science.aaw5570.

Schägger, H., and Jagow, G. von (1991). Blue native electrophoresis for isolation of membrane protein complexes in enzymatically active form. Anal. Biochem. 199, 223–231.

Serre, V., Rozanska, A., Beinat, M., Chrétien, D., Boddaert, N., Munnich, A., Rötig, A., and Chrzanowska-Lightowlers, Z.M. (2013). Mutations in mitochondrial ribosomal protein MRPL12 leads to growth retardation, neurological deterioration and mitochondrial translation deficiency. Biochim. Biophys. Acta 1832, 1304–1312. https://doi.org/10.1016/j.bbadis.2013.04.014.

Silva, D.D., Tu, Y.-T., Amunts, A., Fontanesi, F., and Barrientos, A. (2015). Mitochondrial ribosome assembly in health and disease. Cell Cycle 14, 2226–2250. https://doi.org/10.1080/15384101.2015.1053672.

Tobiasson, V., Gahura, O., Aibara, S., Baradaran, R., Zíková, A., and Amunts, A. (2020). Interconnected assembly factors regulate the biogenesis of mitoribosomal large subunit. Biorxiv 2020.06.28.176446. https://doi.org/10.1101/2020.06.28.176446.

Waltz, F., Soufari, H., Bochler, A., Giegé, P., and Hashem, Y. (2020). Cryo-EM structure of the RNA-rich plant mitochondrial ribosome. Nat Plants 6, 377–383. https://doi.org/10.1038/s41477-020-0631-5.

Westerman, B.A., Poutsma, A., Steegers, E.A.P., and Oudejans, C.B.M. (2004). C2360, a nuclear protein expressed in human proliferative cytotrophoblasts, is a representative member of a novel protein family with a conserved coiled coil–helix–coiled coil–helix domain. Genomics 83, 1094–1104. https://doi.org/10.1016/j.ygeno.2003.12.006.

Zeng, R., Smith, E., and Barrientos, A. (2018). Yeast Mitoribosome Large Subunit Assembly Proceeds by Hierarchical Incorporation of Protein Clusters and Modules on the Inner Membrane. Cell Metabolism 27, 645–656.e7. https://doi.org/10.1016/j.cmet.2018.01.012.

