## Supplementary figures and tables for "Supernumerary proteins of the human mitochondrial ribosomal small subunit are integral for assembly and translation"

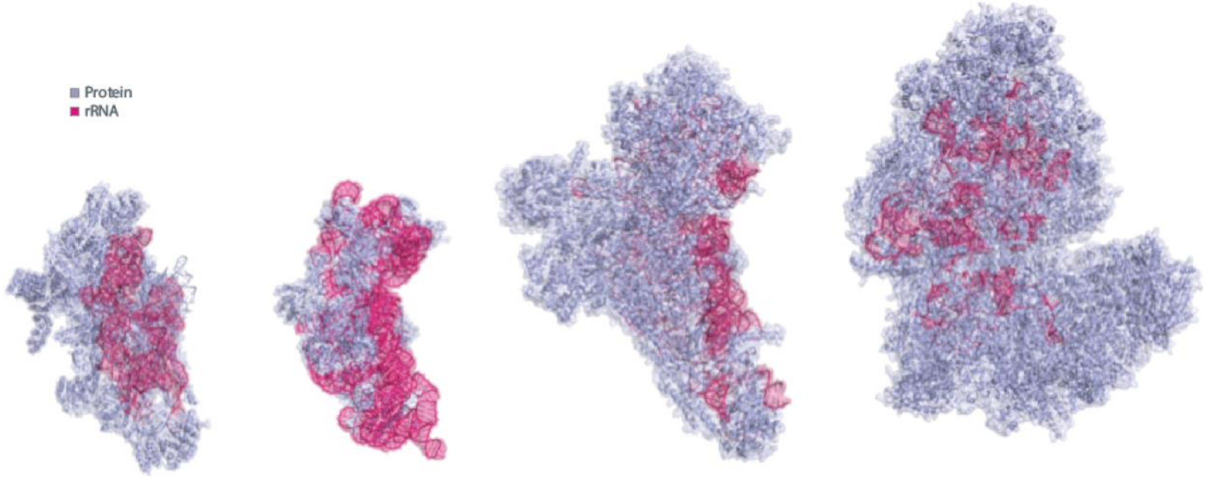


**Supplementary Figure S1 Comparison of *protein-to-RNA composition in the mtSSU of (mitochondrial) ribosomes.***

From left to right: 28S mammalian mitoribosome, 30S bacterial ribosome, 40S yeast mitoribosome, 78S fungi *Neurospora crassa* (full ribosome).

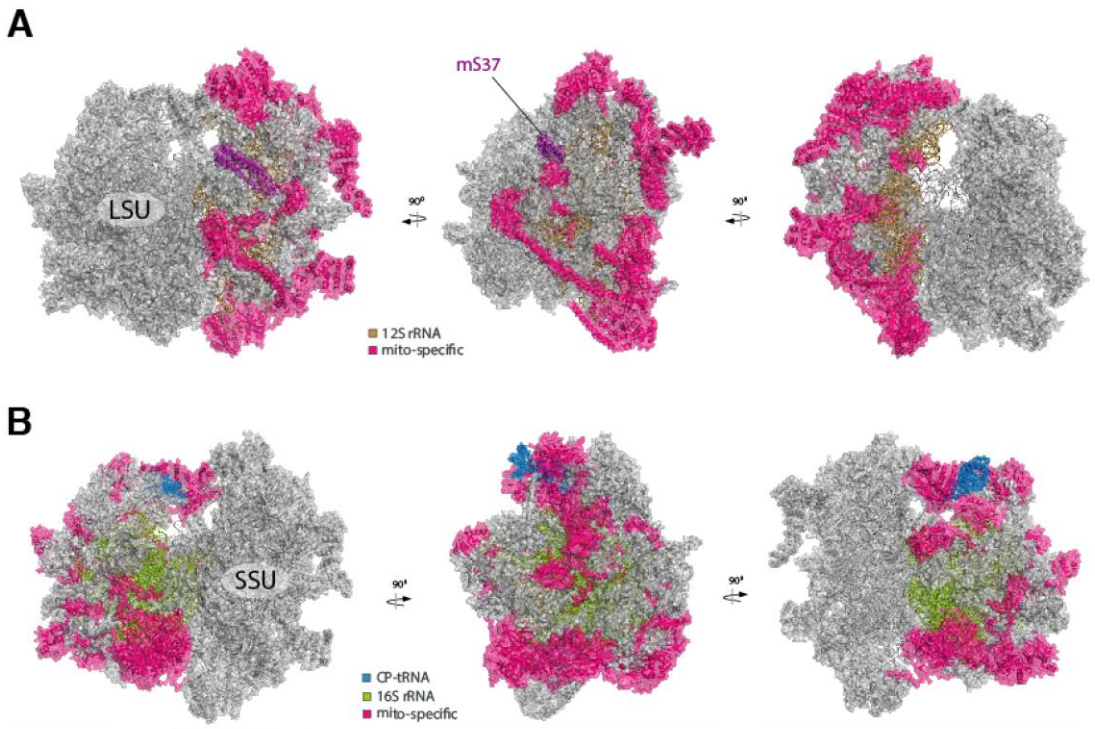


**Supplementary Figure S2** ***Mammalian mtSSU (A) and mtLSU (B).***

(**A**) Location of snMRPs within the mtSSU.
(**B**) Location of snMRPs within the mtLSU.

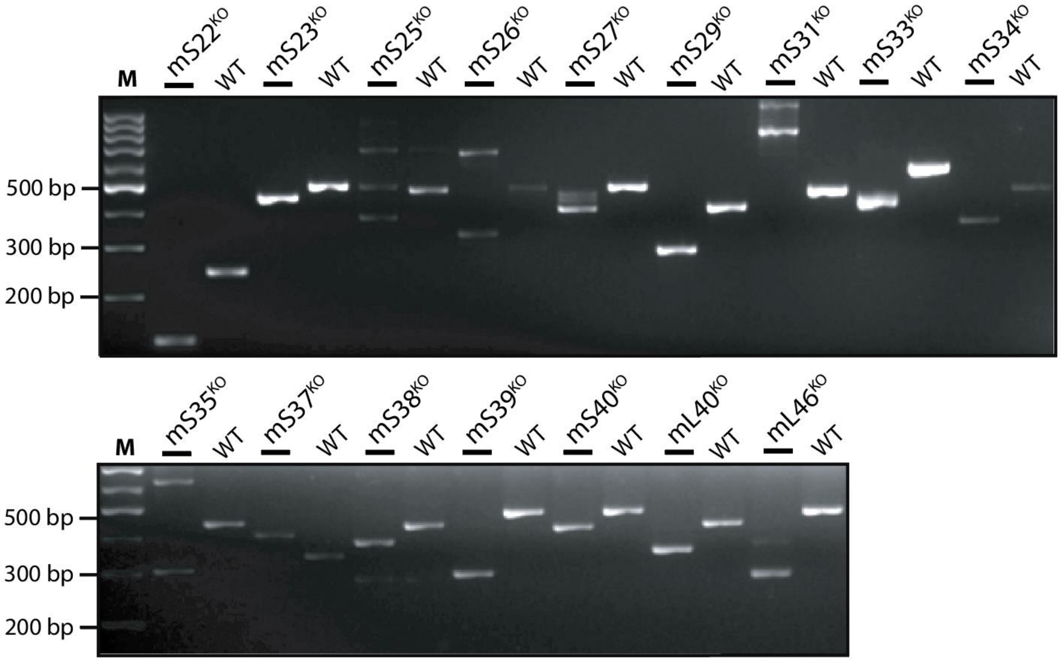


**Figure S3 (related to Figure 1)** ***Genomic DNA amplification of CRISPR-edited genetic loci.*** Primer sequences are listed in Table S4.

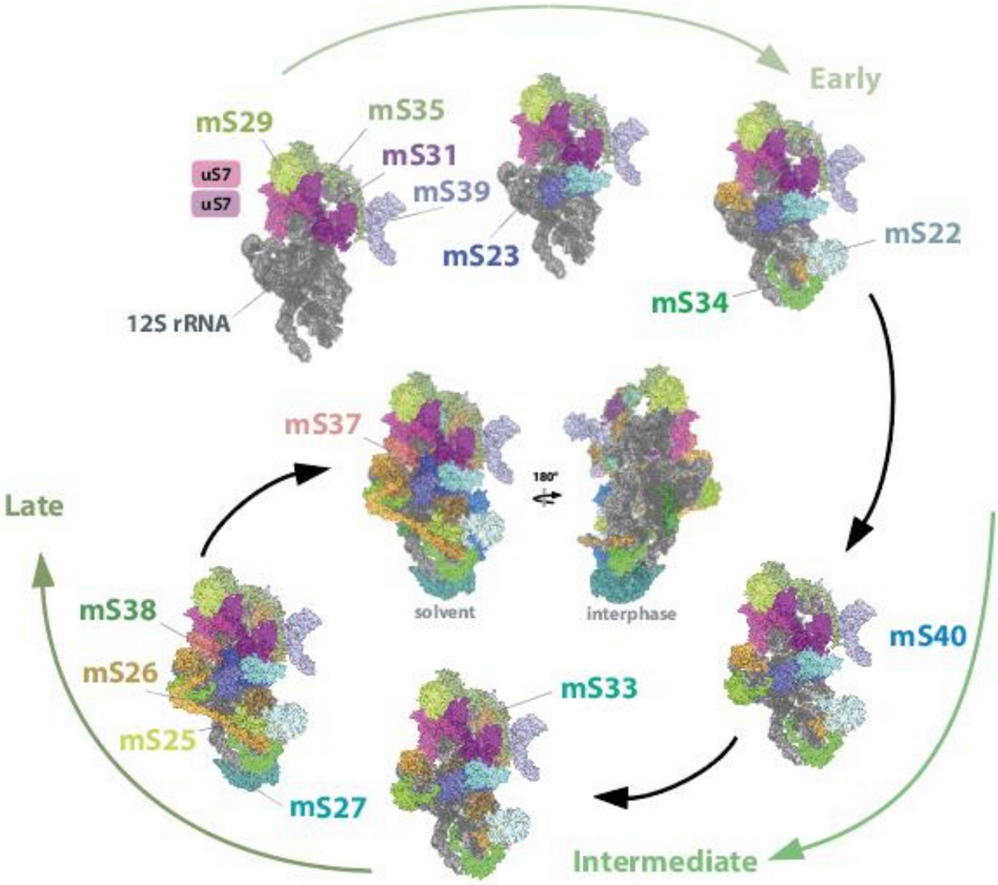


**Figure S4** Representation of mtSSU assembly pathway with snMRPs highlighted (adapted from (Bogenhagen et al., 2018; Ferrari et al., 2020; Hilander et al., 2021)).

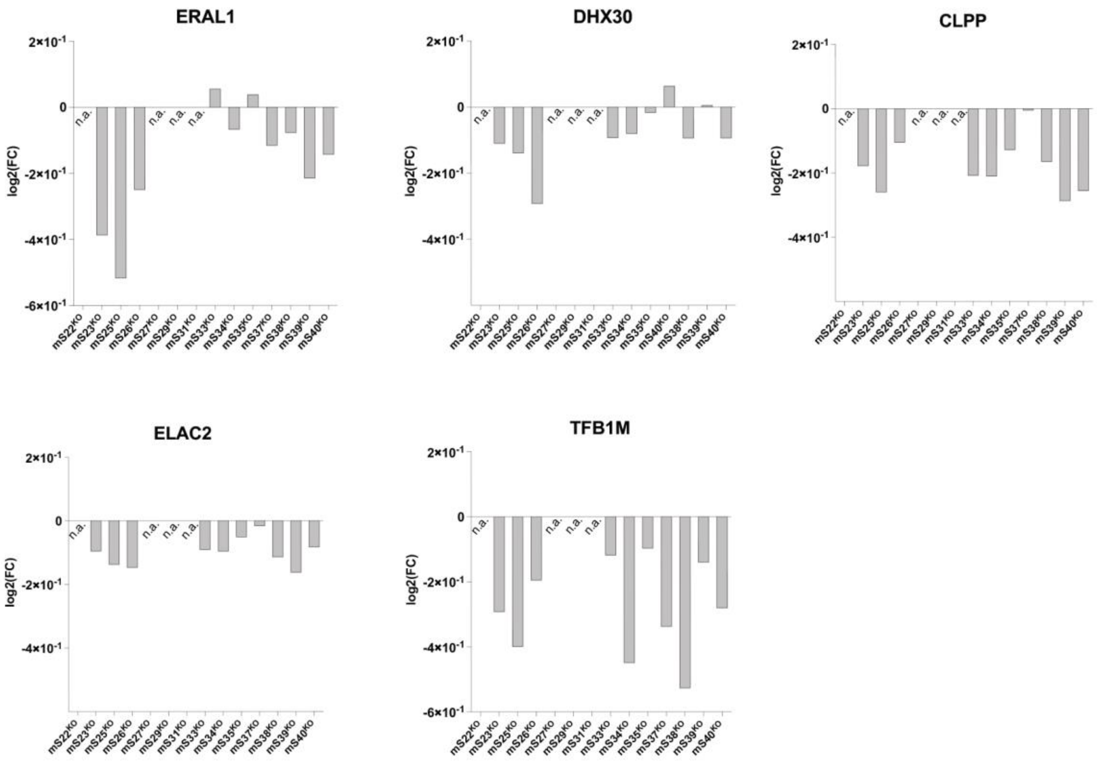


**Figure S5 (related to Figure 3) Ribosome assembly and modifying factors levels from proteomic data.** Values represent mean of quadruplicate measurements normalised to control.

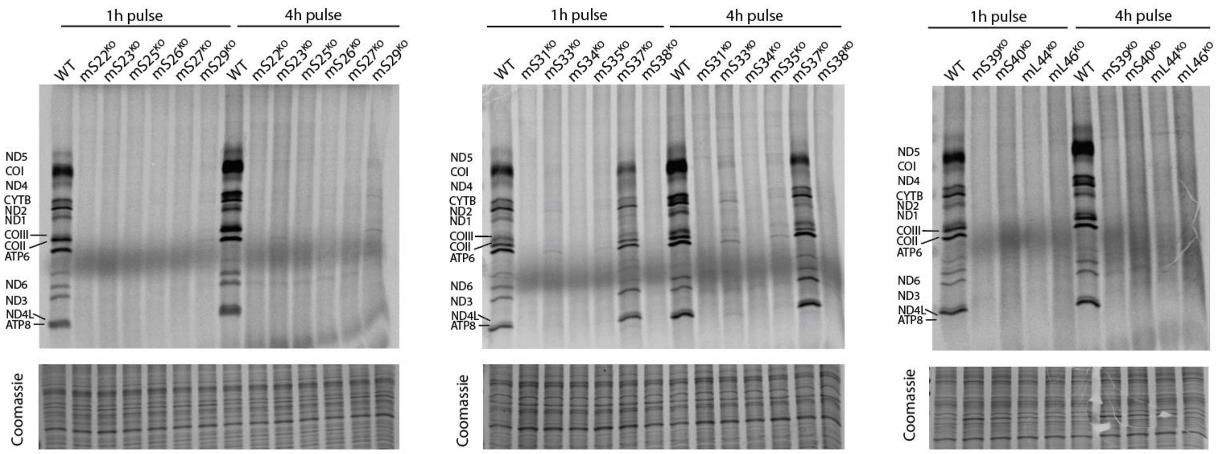


**Figure S6 (related to Figure 4) Extended $^{35}$S-metabolic pulse labelling.**

### *Table S1. Mitochondrial disease-associated mutations in mitoribosomal proteins.*

**mtSSU**

| Subunit / Gene | References |
|---|---|
| uS2m / MRPS2 | (Gardeitchik et al., 2018) |
| uS7m / MRPS7 | (Menezes et al., 2015) |
| uS14m / MRPS14 | (Jackson et al., 2019) |
| uS16m / MRPS16 | (Miller et al., 2004) |
| mS22 / MRPS22* | (Saada et al., 2007) |
| mS23 / MRPS23* | (Kohda et al., 2016) |
| mS25 / MRPS25* | (Bugiardini et al., 2019) |
| bS1m / MRPS28 | (Pulman et al., 2018) |
| mS34 / MRPS34* | (Lake et al., 2017) |
| mS39 / MRPS39* | (Borna et al., 2019) |

* supernumerary subunits

**mtLSU**

| Subunit / Gene | References |
|---|---|
| uL3m / MRPL3 | (Galmiche et al., 2011) |
| bL12m / MRPL12 | (Serre et al., 2013) |
| uL24m / MRPL24 | (Nottia et al., 2020) |
| mL44 / MRPL44* | (Carroll et al., 2013) |

* supernumerary subunits

### *Table S2. gRNAs for CRISPRi-mediated knockout.*

| **20bp crRNAs** | Sequence (5`-3`) |
|---|---|
| mS22 guide 1 | cgcccggagaactcctcaag |
| mS22 guide 2 | tcggagctgaaccggcggcg |
| mS23 guide 1 | ccgagagaagatgctcccta |
| mS23 guide 2 | cggtagcagctagtcacgct |
| mS25 guide 1 | agtgaattacaacacgcatg |
| mS25 guide 2 | gttgccgccatgcccatgaa |
| mS26 guide 1 | atggccgagtctcctcactg |
| mS26 guide 2 | gcgagtgaacatgccgcccg |
| mS27 guide 1 | agatggctgcctccatagtg |
| mS27 guide 2 | ggttacagctcgacctctcg |
| mS28 guide 1 | ccgggctccaccttctgtag |
| mS28 guide 2 | tcagtgcctacaccccgaaa |
| mS29 guide 1 | gggaaaagatgtgtgcaaca |
| mS29 guide 2 | acagagtgttcaggtcagtc |
| mS31 guide 1 | atgctactcactgttcggca |
| mS31 guide 2 | tagcccaccggagcgcaaag |
| mS33 guide 1 | gggcactgagacgagacatg |
| mS33 guide 2 | acactgcactgatccatgag |
| mS34 guide 1 | tggactacgagaccttgacg |
| mS34 guide 2 | aggacttgcgcgtgaccagg |
| mS35 guide 1 | caagcggcgttcagtgtcag |
| mS35 guide 2 | gctaggtgtcgggaccggag |
| mS36 guide 1 | cgggactccagtgatcgccg |
| mS36 guide 2 | tttccgcatcttgggcggta |
| mS37 guide 1 | gatgcaagtcgcctctggag |
| mS37 guide 2 | gcttcctcgattgtgccgcg |
| mS38 guide 1 | gcggcggccacaggtcccag |
| mS38 guide 2 | aggaacggccctcaacagct |
| mS39 guide 1 | gcgtgcctcgaccttcagga |
| mS39 guide 2 | ataggactaaggtgactccg |
| mS40 guide 1 | ctgggcgtacgtcaagatgg |
| mS40 guide 2 | cctgaactctgtgagaacct |
| mL44 guide 1 | cattgcacgagagaaaacga |
| mL44 guide 2 | aaccggagggaccagcttgg |
| mL46 guide 1 | cgttctcccacaatgcaccg |
| mL46 guide 2 | aacggcttggagactacagg |

***Table S3. Oligos for gRNA preparation.***

| Use | Sequence (5`-3`) |
|---|---|
| U6promFw | GAGGGCCTATTTCCCATGATTC |
| U6promRv | GGTGTTTCGTCCTTTCCAC |
| 5pTailedU6promFw | GTAAAACGACGGCCAGTGagggcctatttcccatgattc |
| Term80bpFw | gttttagagctaGAAAtagcaag |
| TermRv80bp | AAAAAAAgcaccgactcggtgccactttttcaagttgataacggactagccttattttaacttgctaTTTCtagctctaaaac |
| 3pTailedTerm80bpRv | AGGAAACAGCTATGACCATGAAAAAAAgcaccgactcggtgccac |
| 1_aggc_Fw | actgaattcggatcctcGAGCGTCTCACCCTGTAAAACGACGGCCAGT |
| 1_aggc_Rv | catgcggccgcgtcgacagatctCGTCTCACATGAGGAAACAGCTATGACCATG |

***Table S4. Screening and sequencing primers.***

| Primers | Sequence (5`-3`) |
|---|---|
| mS22_seqF | aatccctcccaaccacttcc |
| mS22_seqR | cccagcgaaagtccggaa |
| mS23_seqF | gggagaggcagctgcaataat |
| mS23_seqR | ttttggctcggctatcgagt |
| mS25_seqF | gcctcagtctggacctctg |
| mS25_seqR | tacaagtcccagagtgctcc |
| mS26_seqF | ggtcgccgcttcggtt |
| mS26_seqR | caccttcctctgcacctcg |
| mS27_seqF | taggctaaagccgcggatac |
| mS27_seqR | tctaagacccagcaggtggta |
| mS29_seqF | cactagccttttgtgtttcgt |
| mS29_seqR | tccatggtaaattctcaagcaca |
| mS31_seqF | tctaagacccagcaggtggta |
| mS31_seqR | cctttgctgaactctggcga |
| mS33_seqF | acctgggctctattataagaacaa |
| mS33_seqR | gcaaaggaggcaatacagca |
| mS34_seqF | ctgccacagccaggacttg |
| mS34_seqR | tcagcgtcggagtctgagat |
| mS35_seqF | gggggaatcttcctgcacat |
| mS35_seqR | cagacccacgtccatggttt |
| mS36_seqF | acacgtgggtggatcctagt |
| mS36_seqR | tccaacaggctcaaagtccc |
| mS37_seqF | gagcagtcggagtcaggac |
| mS37_seqR | cccaaatgaatgaaagggcct |
| mS38_seqF | ttctgggacctttcggtgcg |
| mS38_seqR | tccctctgtgagcatcgga |
| mS39_seqF | attcctccgaggcaaatcgg |
| mS39_seqR | tgcgttgcgaatcctatttca |
| mS40_seqF | tgcgatctaagagtcgtagtgac |
| mS40_seqR | cgtttagtcctcggctcgg |
| mL44_seqF | cctgccctctctcagtcg |
| mL44_seqR | tgtctccatcgcaaacttcc |
| mL46_seqF | gttctagcagttccgggact |
| mL46_seqR | acggccaatgaaaagaaacca |

### *Table S5. Antibodies used in this study.*

| Reagent | Source /Identifier | Dilution |
|---|---|---|
| MRPS22/mS22 | Proteintech 10984-1-AP | 1:1000 |
| MRPS23/mS23 | Proteintech 18345-1-AP | 1:1000 |
| MRPS25/mS25 | Proteintech 15277-1-AP | 1:1000 |
| MRPS26/mS26 | Proteintech 15989-1-AP | 1:1000 |
| MRPS27/mS27 | Proteintech 17280-1-AP | 1:1000 |
| MRPS29/mS29 | Proteintech 10276-1-AP | 1:1000 |
| MRPS31/mS31 | Proteintech 16288-1-AP | 1:1000 |
| MRPS34/mS34 | Proteintech 15166-1-AP | 1:1000 |
| MRPS35/mS35 | Proteintech 16457-1-AP | 1:1000 |
| MRPS37/mS37 | Proteintech 11728-1-AP | 1:500 |
| PTCD3/MRPS39/mS39 | Proteintech 25158-1-AP | 1:1000 |
| MRPS18B/MRPS40/mS40 | Proteintech 16139-1-AP | 1:1000 |
| MRPL44/mL44 | Proteintech 16394-1-AP | 1:1000 |
| MRPL46/mL46 | Proteintech 16611-1-AP | 1:1000 |
| MRPL11/uL11m | Proteintech 15543-1-AP | 1:20000 |
| α-tubulin | Sigma-Aldrich T5168 | |
| NDUFA9/Complex I | MitoSciences D0314 | |
| SDHA | Abcam ab14715 | 1:10000 |
| CytB/Complex III | Proteintech 55090-1-AP | 1:1000 |
| MTCO1/COXI/Complex IV | ThermoFisher Scientific 1D6E1A8 | 1:500 |
| MTCO2/COXII/Complex IV | Proteintech 55070-1-AP | 1:1000 |
| ATP5B/Complex V | Proteintech 17247-1-AP | 1:1000 |
